# MvPPT: a highly efficient and sensitive pathogenicity prediction tool for missense variants

**DOI:** 10.1101/2022.01.05.475156

**Authors:** Shi-Yuan Tong, Ke Fan, Zai-Wei Zhou, Lin-Yun Liu, Shu-Qing Zhang, Yinghui Fu, Guang-Zhong Wang, Ying Zhu, Yong-Chun Yu

## Abstract

Next generation sequencing technologies both boost the discovery of variants in the human genome and exacerbate the challenges of pathogenic variant identification. In this study, we developed mvPPT (Pathogenicity Prediction Tool for missense variants), a highly sensitive and accurate missense variant classifier based on gradient boosting. MvPPT adopts high-confidence training sets with a wide spectrum of variant profiles, and extracts three categories of features, including scores from existing prediction tools, allele, amino acid and genotype frequencies, and genomic context. Compared with established predictors, mvPPT achieved superior performance in all test sets, regardless of data source. In addition, our study also provides guidance for training set and feature selection strategies, as well as reveals highly relevant features, which may further provide biological insights of variant pathogenicity.

## Introduction

Whole exome sequencing (WES) and whole-genome sequencing (WGS) enable large-scale parallel assessment of genetic variants and has been increasingly adopted in clinical diagnosis, which makes interpreting the effect of the identified variants a serious challenge (1,2). Unlike synonymous and loss-of-fUnction variants for which the impact on the protein can be relatively easy to predict, missense variants that often lead to inconclusive genomic outcomes remains a major challenge in pathogenicity interpretation. Compared with the reference genome, a human exome on average contains around 20,000 single-nucleotide variants and approximately half are missense variants (3,4). Nevertheless, the effects of most missense variants on proteins are unclear, as experimental validation of large numbers of variants is limited by efficiency and cost. To address these limitations, many computational tools have been developed to predict the potential impact of variants (5–21). Early prediction models compute deleterious scores based on single property of variants, such as evolutionary conservation (8,10,15) and protein structure/function (16,17), and recent ensemble methods achieve higher classification accuracy by integrating information from individual predictors (5–7,9,11–13,20). Although these existing tools have made significant contributions to the prediction of the hazard of genetic variants, the sensitivity of prediction still needs to be improved when assessing the pathogenicity of massive variants in clinical scenarios.

While the existed tools provide positive predictive power, their prediction results are often inconsistent with each other (6,18). It is believed that the predictive power of current ensemble methods is hampered by the lack of appropriate training data and incomplete features (7,8,22). For training set inclusion, the widely-adopted strategies to create training sets include using variants from disease databases only (7,9), or using variants from both disease and population databases to balance the ratio of benign and pathogenic variants in the dataset (5,6). However, there is no conclusion on which strategy results in the best performance. Specifically, existing ensemble tools mostly train machine learning models on variants with known labels in disease databases such as ClinVar (23) and/or Human Gene Mutation Database (HGMD) (24). However, variants in disease databases may cluster around well-described disease genes, i.e., the more a gene has been studied, the more variants on this gene are likely to be discovered. Unfortunately, it is not clear yet whether the clustering of variants on certain genes introduce bias in the prediction of variant pathogenicity. Moreover, since each resource maintains different variant inclusion criteria, vast variants in ClinVar and HGMD databases were labeled with conflict or even opposite clinical significance (25,26), which might attenuate the prediction accuracy of the computational tools. To further expand the training data, some existing tools include sequence variants from population databases, such as Exome Sequencing Project (ESP) (6,11,27). Most of these tools consider the sequence variants in general populations above a certain allele frequency (AF) as benign, however, how to choose a proper AF threshold for defining neutral training variants remains a question.

The common features adopted by most of the present ensemble models are scores computed by individual predictors based on amino acid or nucleotide conservation, and biochemical properties of the amino acid substitutions (5,6,9,12,13). While these scores are proven to be highly relevant to variant deleteriousness, other features linked to variant pathogenicity have been shown strong correlation with human disease. For example, AF has been used as an important criterion in deleterious selection in practice for a long time, but was rarely considered in an ensemble model. ClinPred add allele frequencies as input features for the first time, and is shown to be more effective than many other ensemble machines (7). Similarly, genotype frequency (GF) and amino acid frequency (AAF) contain hints of natural selection, which may provide extra information for pathogenicity inference. Additionally, recent studies have shown that intolerance to variation is a strong predictor of human disease relevance, emphasizing the role of genomic context in variant pathogenicity prediction (28–30).

As the algorithm is the “brain” of a machine learning model, the efficiency of a model is largely dependent on algorithm selection. Varieties of machine learning approaches, such as logistic regression (9,13), support vector machine (SVM) (9), random forest (6,13), and boosting algorithms (5,7) have been implemented in variant classification. In general, tree-based approaches achieve higher accuracy and precision according to prior studies(5–7,19), however, few studies have systematically evaluated the effects of different algorithms on pathogenic variant prioritization. LightGBM is a gradient boosting framework that uses tree-based learning algo1rithms (31). Unlike random forests where the component trees are trained independently, in gradient boosting, trees are built in a stepwise manner, where each successive tree is optimized on the residuals of the prediction of the preceding tree. In previous studies, it has been demonstrated that compared with other gradient boosting frameworks such as XGBoost and Catboost, LightGBM converges on a solution that generalizes better (32).

Considering the aforementioned observations, we introduce mvPPT, a novel gradient boosting machine for missense variant pathogenicity prediction. By selecting 62 features, including scores from individual predictors, AF/GF/AAF, and genomic context information, and adopting a high-confidence variant training set, mvPPT demonstrated a best-to-date performance in variant pathogenicity prediction, paving its way in molecular diagnosis and clinical scenario applications. MvPPT and pre-computed scores of missense variants in the human exome can be accessed through http://www.mvppt.club.

## Results

### MvPPT design

The performance of a machine learning model is mainly determined by the algorithm, the features, and the training set it used. Therefore, we designed mvPPT by carefully selection of the algorithm, features and the training set (**Figure 1**).

**Figure 1.**
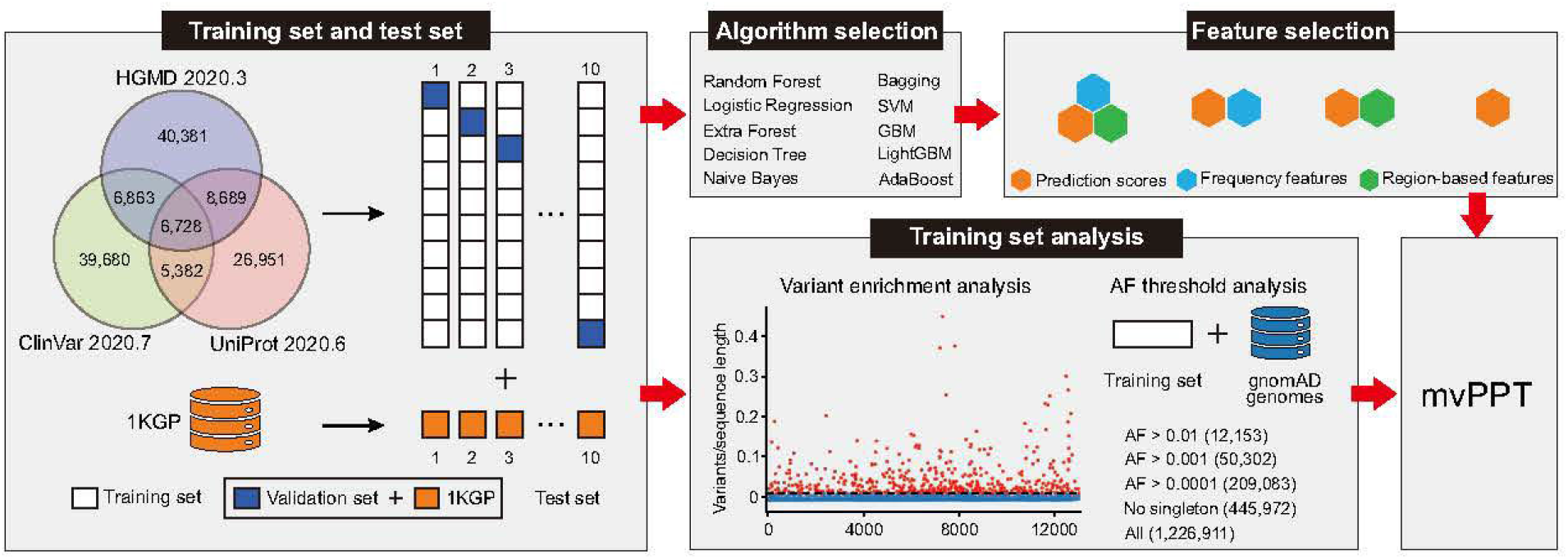
The detailed mvPPT construction process. MvPPT workflow. High-confidence variant sets were extracted from ClinVar, HGMD, and Uniprot. Models were trained using LightGBM with parameters tuned by Bayesian global optimization. ten-fold cross-validation is carried out to verify the effectiveness of the prediction model. MvPPT was built after algorithm selection, feature selection, and training set analysis.

### Data sets for algorithm and feature selection

In order to obtain high-confidence training data, we enrolled variants from currently biggest disease databases: ClinVar, HGMD, and Uniprot (33). Additionally, in order to mimic real world conditions where there are more benign variants than pathogenic variants, we created a benign variant set including population sequence variants from the 1000 Genomes Project (1KGP) (34) removing singletons (**Figure 1**).

#### Algorithm

We first benchmarked the performance of ten commonly used algorithms, including SVM, naive bayes, logistic regression, decision tree, random forests, extra forests, gradient boosting machine (GBM), AdaBoost, LightGBM, and bagging on the data test set mentioned above. To achieve a fair comparison, the default parameters were used for all algorithms. The performance of each algorithm was evaluated using the area under the receiver operating characteristic curve (AUROC) and the area under the precision-recall curve (AUPRC). AUROC and AUPRC range from 0-1, and the closer the metrics are to 1, the better the models are. We found that the performance of the ensemble learning (random forests, extra forest, GBM, AdaBoost, LightGBM, and bagging) achieved the higher AUROC and AUPRC (**Figure 2**). Among them, LightGBM has the highest AUROC (0.970 ± 0.001) and AUPRC (0.952 ± 0.002, *P*-value = 0.0020 vs second-best, Wilcoxon matched-pairs signed rank test) (**Figure 2**).

**Figure 2.**
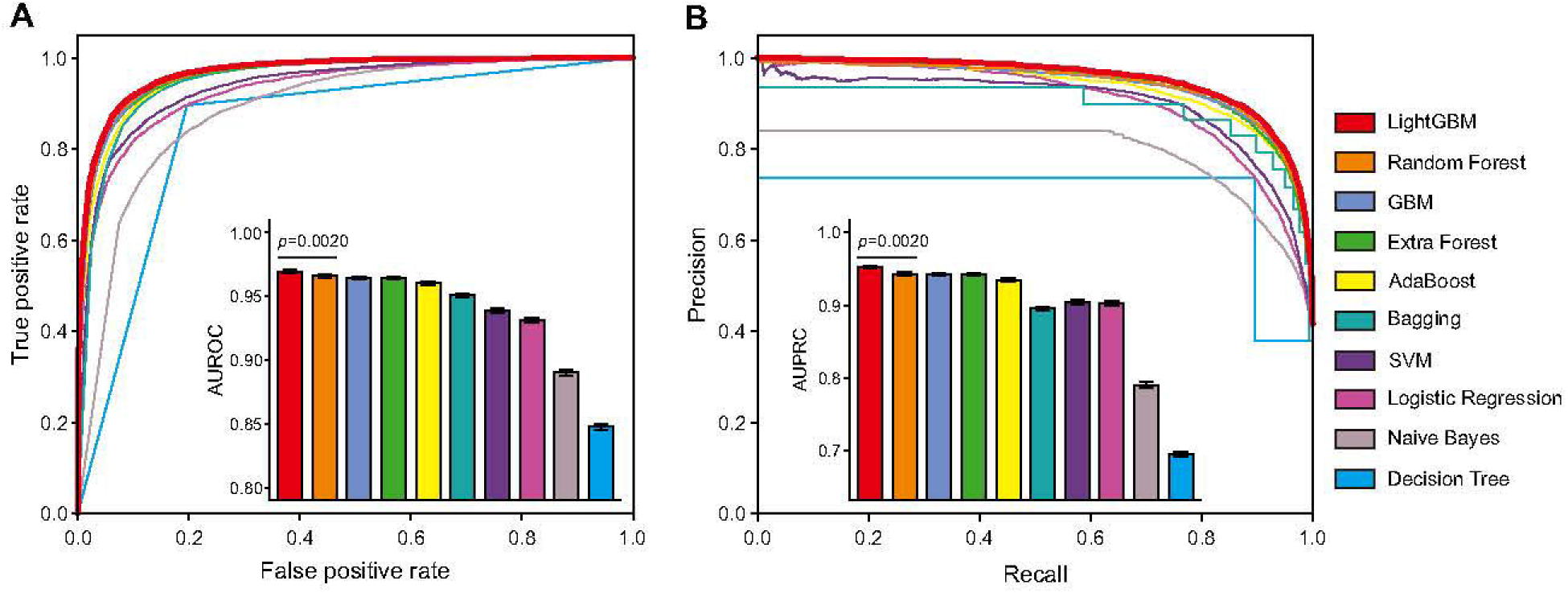
Comparing the performance of different algorithms. Performance assessment of models using different algorithms. (**A**) The operating characteristic curve (ROC) and (**B**) the precision-recall curve (PRC) for models trained on different algorithms were plotted. Below: barplot quantifications of the AUROC and the AUPRC by using ten-fold cross-validation. *P*-value was calculated by Wilcoxon matched-pairs signed rank test.

#### Features

To obtain a proper feature space, we extracted three types of features, including category A: pathogenicity likelihood scores generated by 7 non-machine learning-based component tools, including MutationAssessor (15), phyloP (35), GERP++RS (36), phastCons (37), PROVEAN (38), SiPhy (39), and SIFT (10), category B: AF, AAF, and GF of variants estimated from 125,748 exomes in gnomAD (version 2.1.1) (28), and category C: region/gene-based information obtained from GeVIR (29), VIRLoF (29), oe_mis_upper (from gnomAD), HIP (40), CCRs (41), Interpro domain (42), and amino acid before and after mutation (**Table 1; Figure 1**). To test if each category of features represents distinct discriminative information, we evaluated the performance of models with the following combinations of feature categories: A, A+B, A+C, and A+B+C (**Table 1**). Overall, adding category B and/or category C greatly boost the performance, compared with the model including only category A, which is the case of most previous ensemble machines. Specifically, the model including A+B+C achieved the highest performance (**Figures 3A, B**). As pathogenic variants in databases may be enriched with low-frequency variants, we also assessed the performance of different models on rare variants. When all the variants in test set were rare (AF = 0, based on gnomAD exomes), the main contributions come from category A and category C, and adding category B shows few but slightly positive impacts (**Figures 3C, D**). Altogether, both category B and category C improved forecasting accuracy and were thus included in mvPPT.

**Table 1.** Features associated with missense variants mined in this work.

| Category | Features | Definition |
| --- | --- | --- |
| A | MutationAssessor, SIFT, PROVEAN, GERP++RS, phyloP100way_vertibrate, phyloP30way_mammalian, phyloP17way_primate, phastCons100way_vertibrate, phastCons30way_mammalian, phastCons17way_primate, SiPhy_29way_logOdds | variant prediction scores from different component tools |
| B | gnomAD_WtFs<sup>a</sup>, gnomAD_AFs<sup>b</sup>, gnomAD_HetFs<sup>c</sup>, gnomAD_HomFs<sup>d</sup>, gnomAD_AAFs<sup>e</sup> | AAFs, AFs, and GFs of each variant from the gnomAD exomes database |
| C | GeVIR, VIRLoF, HIP, CCRs, Interpro_domain, oe_mis_upper, <sup>f</sup>RefAA_A, RefAA_C, RefAA_D, RefAA_E, RefAA_F, RefAA_G, RefAA_H, RefAA_I, RefAA_K, RefAA_L, RefAA_M, RefAA_N, RefAA_P, RefAA_Q, RefAA_R, RefAA_S, RefAA_T, RefAA_V, RefAA_W, RefAA_Y, <sup>g</sup>AltAA_A, AltAA_C, AltAA_D, AltAA_E, AltAA_F, AltAA_G, AltAA_H, AltAA_I, AltAA_K, AltAA_L, AltAA_M, | Region/gene-based scores |

*Note:* <sup>a</sup>Wildtype frequencies estimated from all exomes in gnomAD. <sup>b</sup>Allele frequencies estimated from all exomes in gnomAD. <sup>c</sup>Heterozygous frequencies estimated from all exomes in gnomAD. <sup>d</sup>Homozygous frequencies estimated from all exomes in gnomAD. <sup>e</sup>Amino acid frequencies estimated from all exomes in gnomAD. <sup>f</sup>Reference amino acid. <sup>g</sup>Alternate amino acid.

**Figure 3.**
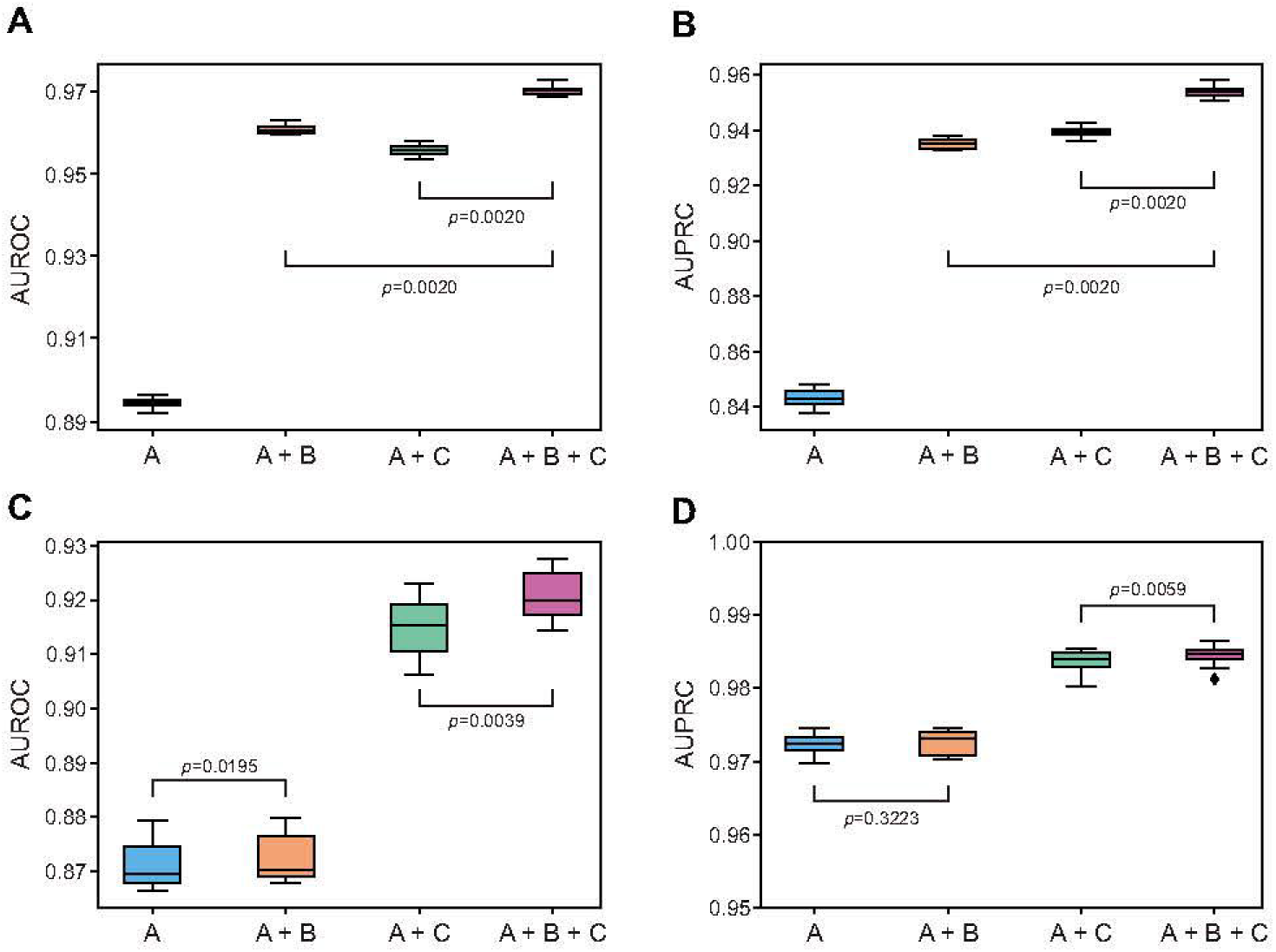
The prediction power of features by category. (**A**) The AUROC and (**B**) AUPRC obtained from models trained on each combination of categories were plotted. (**C**) AUROC and (**D**) AUPRC evaluating the performance of models trained on different combinations of categories when all the variants in the test set were rare (AF = 0, based on gnomAD exomes). AUROC and AUPRC from ten-fold cross-validation were plotted. *P*-value was calculated by Wilcoxon matched-pairs signed rank test.

As population variants tend to have higher AFs than pathogenic variants, modes including AFs as features may perform better in test sets including population variants. Therefore, we excluded 1KGP variants from the test set and re-conducted the comparisons models displayed similar performance on test sets with or without population variants (**Figure S1**).

#### Training set

To obtain a proper training set, we first evaluated the biases of variant collection in different databases, and then compared the performance of models trained on training sets generated using different strategies.

We observed that genomic locations of variants recorded in disease databases are likely to be biased by interests of the research field, i.e., variants in the databases are likely to be enriched in “hotspots” of the human genome. To evaluate the enrichment pattern of variants from different databases on genome, we calculated the ratio between the number of missense variants in each gene and the length of the gene’s protein-coding sequence (VPR). We found that VPR in disease databases is much variable than that in gnomAD, with coefficient of variation (C.V) 0.815% for gnomAD and 2.35% for disease databases (**Figures 4A, C**). Outlier detection based on interquartile range (IQR) detected 1.12% of the genes as outliers (VPR > mean + 1.5 * IQR) in gnomAD, but 13.48% of the genes as outliers in disease databases (**Figures 4B-D and Table S1**). Likewise, when plotting the number of variants against the length of coding sequence, we observed a significant positive correlation for variants in gnomAD, but not for variants in the pathogenic databases (**Figures 4E, F**). Gene ontology (GO) enrichment analysis revealed that outlier genes in gnomAD are enriched in pathways associated with immune response (43–45), which are known to be hotspots of positive selection. In contrast, top enriched pathways of outlier genes in pathogenic databases include gland development (*p*.adjust = 1.33E-12), regulation of body fluid levels (*p*.adjust = 8.61E-12), and response to inorganic substance (*p*.adjust = 9.70E-11) (**Figures 4G, H**), reflecting that different variant enrichment patterns are there in the disease and population databases.

**Figure 4.**
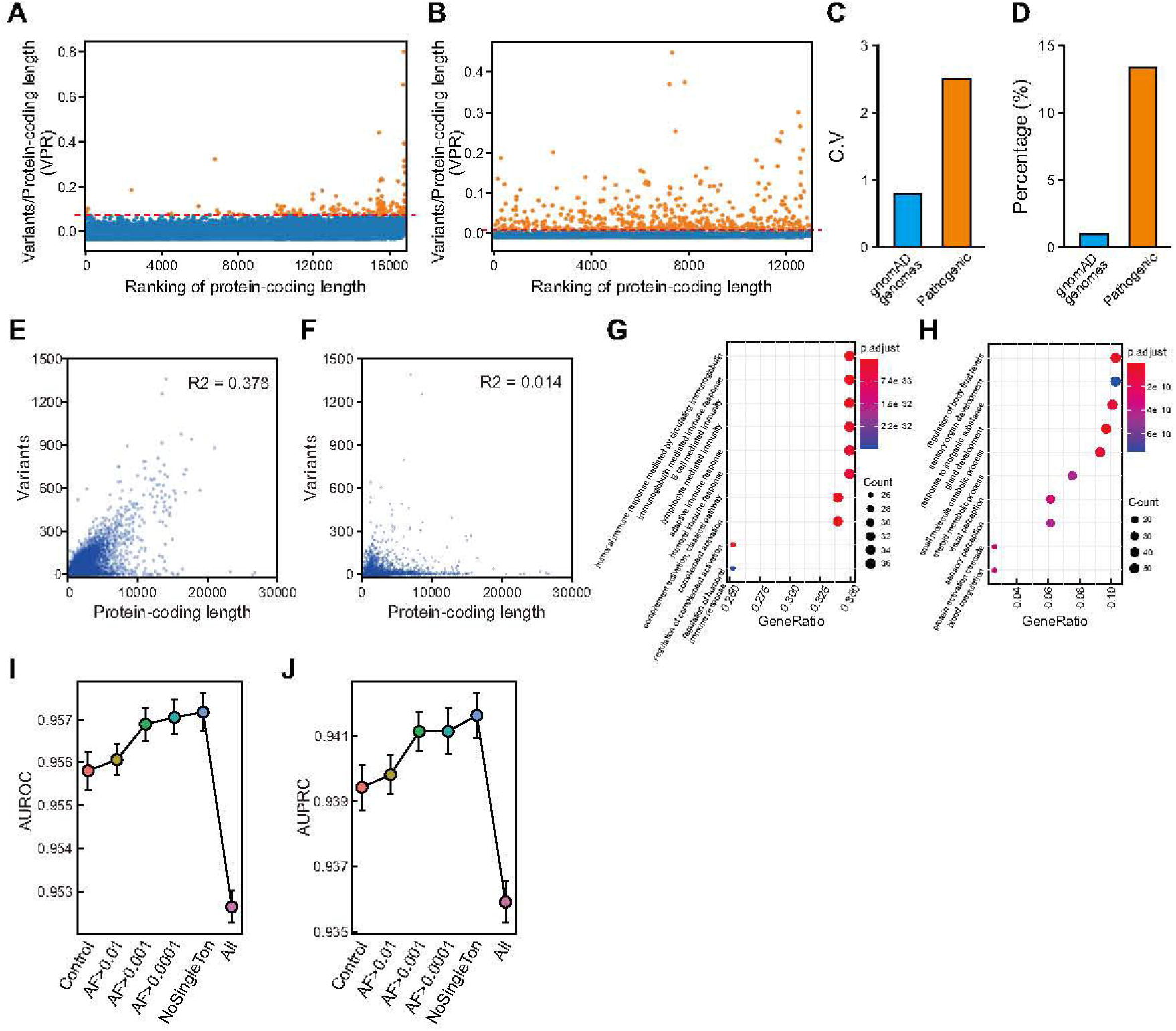
The enrichment pattern of variants from different databases and AF selection. Ratio between the number of missense variants in each gene and the length of the gene’s protein-coding sequence length (VPR) was calculated. (**A**) After been centered, VPR in gnomAD genomes was plotted against the ranking of protein-coding sequence length. (**B**) Same as (**A**) but for pathogenic variants. (**C**) The C.V. for VPR in gnomAD genomes and in disease databases. (**D**) The percentage of outlier genes related to gnomAD genomes and in disease databases. (**E**, **F**) Variants number profile across all genes in gnomAD genomes (**E**) and in disease databases (**F**). R^2^ corresponds to the coefficient of determination. (**G**, **H**) Bubble chart showing GO enrichment analysis of variants in gnomAD genomes and in disease databases. (**I**) AUROC and (**J**) AUPRC obtained from models trained on training sets combining gnomAD variants selected according to different AF thresholds were plotted. Control represents the model without incorporating gnomAD variants.

To further test if aggregation of genetic variants on genes impairs the model performance, we down-sampled pathogenic variants on genes with large numbers of pathogenic variants. Specifically, on genes with VPR greater than a setting threshold (0.008, 0.015, 0.040, 0.065, and 0.090), we randomly selected a fixed number of variants and combined them with variants from other genes to form the down-sampled training sets (**Figure S2A**). The down-sampled training sets were then fed to models using different feature combinations (A+B and A+B+C). To avoid similar variant enrichment pattern in the test set, we randomly selected one pathogenic variant from each gene to form a test set. Ten test sets were created in each round of the ten-fold cross-validation. We found that the predictive power of the models reduced with down-sampling (**Figures S2B, C**), which is possibly because the number of variants available for learning is largely reduced with down-sampling. Adding category C slowed down the reduction. But, overall, down-sampling appears to attenuate the performance of the model, and thus is disfavored (**Figures S2B, C**).

Next, we compared the performance of models trained on six training sets generated by different strategies. In our assessment, we found that adding variants from population databases skewed the distribution of benign variant AFs towards zero, making it similar to that of pathogenic variants (**Figure S3**). Including neutral variants from the population database have a positive impact on the model, with adding variants removing singletons displaying the highest performance (**Figures 4I, J**). In contrast, adding the full variants from gnomAD jeopardizes the model performance, possibly due to the inclusion of variants in the population database that are not true benign variants (**Figures 4I, J**). To further investigate whether this improvement is due to correlation of AFs between training and test sets, we divided the original test sets into bins based on AFs (based on gnomAD exomes), and tested the performance of the models on variants within a specific AF range. We found that including population variants removing singleton slightly enhances the performance of the model in most of the bins, especially the low-AF bins, where most of the pathogenic variants located, likely because it expands the training set while excluding unreliable (singleton) samples (**Table S2**). In contrast, the performance of the model drops with AF cutoff increases, probably due to lack of rare benign variants in the training set.

Based on the above analyses, mvPPT was finally trained using LightGBM (tuned by Bayesian optimization) on variants from three disease databases and gnomAD with singleton removed, using all features from category A, B, and C. The correlation among the individual features and features relative importance of these features is shown in **Figure S4** and the description of training set is shown in **Table S3**.

### MvPPT outperforms existing prediction tools

For assessment, we collected variants from VariSNP (46), VKGL (47), DPV (48), DoCM (49), MetaLR/SVM_Test (9), and CAPICE_Test (19), to generate an independent test set. Variants that were used in any training set or in our features’ training data were discarded from test sets, and only variants with comprehensive scores required by all comparators were included. In total, there are 175,144 variants in test set with 168,222 benign variants and 6,922 pathogenic variants (**Table S4**).

Using the new test set, the performance of mvPPT was benchmarked against 15 prediction tools that are widely used and readily implemented, including MVP (12), CAPICE (19), FATHMM-XF (20), REVEL (6), M-CAP (version 1.4) (5), ClinPred (7), ReVe (22), PrimateAI (8), MetaSVM (9), MetaLR (9), MISTIC (13), CADD (version 1.4) (21), PolyPhen-2 HDIV (16), PolyPhen-2 HVAR (16), and VEST (version 4) (11). Among all these tools, mvPPT has the highest AUROC of 0.960 and the highest AUPRC of 0.791 (**Figure 5**). MISTIC has the second-best overall performance, with AUROC of 0.920 and AUPRC of 0.565. Besides, we also calculated other metrics include F1 score, accuracy (ACC), positive predictive value (PPV, also known as precision), true positive rate (TPR, also known as sensitivity), Matthews correlation coefficient (MCC), true negative rate (TNR, also known as specificity), and false positive rate (FPR), log loss value, diagnostic odd ratio (DOR) (**Table 2**). M-CAP has the highest recall of 0.953 (second-best: mvPPT, 0.888), but this comes at the cost of a low precision of 0.078 (best: mvPPT, 0.323). PrimateAI has the highest specificity of 0.925 (second-best: mvPPT, 0.923) and lowest FPR of 0.075 (second-best: mvPPT, 0.077). Here, mvPPT has the highest ACC of 0.922, the highest F1 score of 0.473, the highest DOR of 95.102, the highest MCC value of 0.508, the highest precision of 0.323, as well as the lowest log loss (**Table 2**).

**Figure 5.**
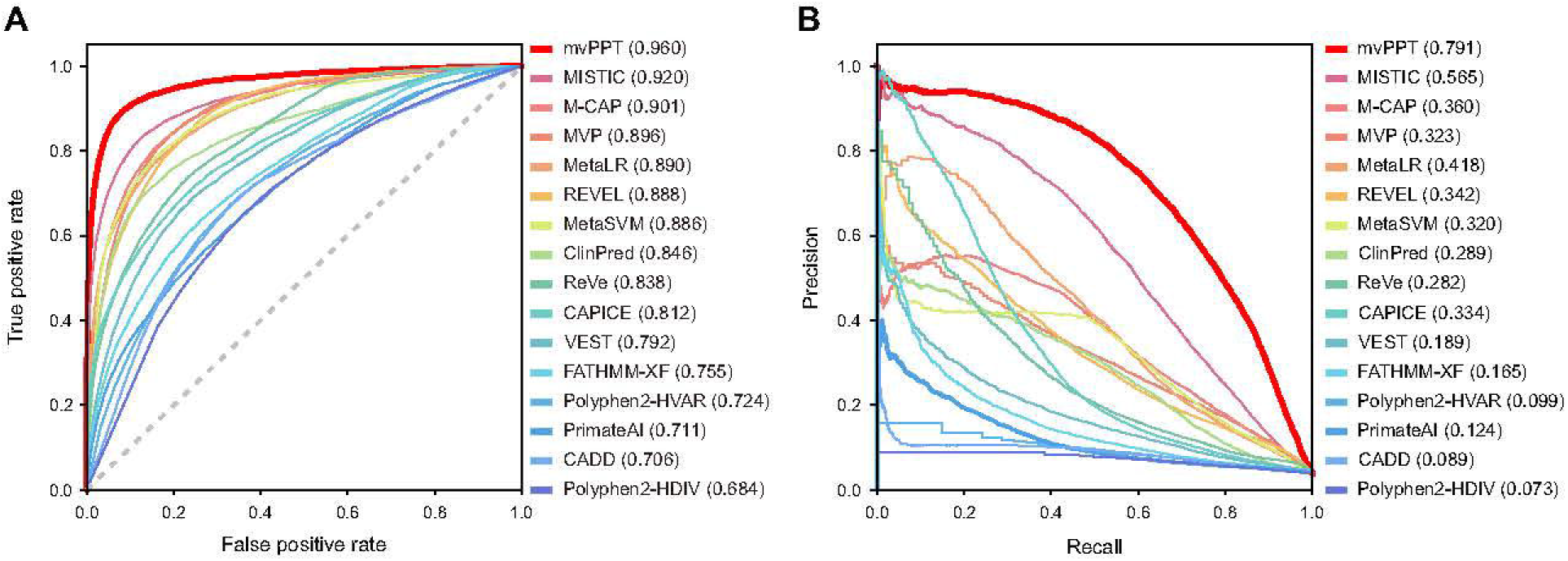
Comparing the performance of mvPPT against established prediction tools. (**A**) AUROC and (**B**) AUPRC of mvPPT and 15 established prediction methods when tested on an independent test set were plotted. The 15 methods included MVP, CAPICE, FATHMM-XF, REVEL, M-CAP, ClinPred, ReVe, PrimateAI, MetaSVM, MetaLR, MISTIC, CADD PolyPhen-2 HDIV, PolyPhen-2 HVAR, and VEST.

**Table 2.** Overview of performance of mvPPT in comparison to other tools in test set.

|  | AUROC<sup>a</sup> | AUPRC<sup>b</sup> | Accuracy | Precision | Recall | F1 score | Log loss | MCC<sup>c</sup> | TNR<sup>d</sup> | FPR<sup>e</sup> | DOR<sup>f</sup> |
| --- | --- | --- | --- | --- | --- | --- | --- | --- | --- | --- | --- |
| mvPPT | 0.960 | 0.719 | 0.922 | 0.323 | 0.888 | 0.473 | 2.697 | 0.508 | 0.923 | 0.077 | 95.102 |
| MISTIC | 0.920 | 0.565 | 0.863 | 0.203 | 0.839 | 0.327 | 4.718 | 0.371 | 0.864 | 0.136 | 33.238 |
| M-CAP | 0.901 | 0.360 | 0.551 | 0.078 | 0.953 | 0.144 | 15.507 | 0.190 | 0.534 | 0.466 | 23.381 |
| MVP | 0.896 | 0.323 | 0.773 | 0.134 | 0.869 | 0.232 | 7.853 | 0.285 | 0.769 | 0.231 | 22.063 |
| MetaLR | 0.890 | 0.418 | 0.830 | 0.160 | 0.782 | 0.266 | 5.889 | 0.303 | 0.831 | 0.169 | 17.726 |
| REVEL | 0.888 | 0.342 | 0.849 | 0.171 | 0.733 | 0.278 | 5.204 | 0.305 | 0.854 | 0.146 | 16.087 |
| MetaSVM | 0.886 | 0.320 | 0.847 | 0.175 | 0.772 | 0.286 | 5.271 | 0.320 | 0.851 | 0.149 | 19.266 |
| ClinPred | 0.846 | 0.289 | 0.683 | 0.096 | 0.829 | 0.171 | 10.942 | 0.208 | 0.677 | 0.323 | 10.157 |
| ReVe | 0.838 | 0.282 | 0.608 | 0.080 | 0.856 | 0.147 | 13.545 | 0.179 | 0.598 | 0.402 | 8.817 |
| CAPICE | 0.812 | 0.334 | 0.733 | 0.101 | 0.731 | 0.178 | 9.224 | 0.200 | 0.733 | 0.267 | 7.457 |
| VEST | 0.792 | 0.189 | 0.634 | 0.079 | 0.780 | 0.144 | 12.638 | 0.163 | 0.628 | 0.372 | 5.997 |
| FATHMM-XF | 0.755 | 0.165 | 0.586 | 0.069 | 0.761 | 0.127 | 14.285 | 0.134 | 0.579 | 0.421 | 4.388 |
| Polyphen2_HVAR | 0.724 | 0.099 | 0.687 | 0.079 | 0.651 | 0.141 | 10.809 | 0.141 | 0.689 | 0.311 | 4.128 |
| PrimateAI | 0.711 | 0.124 | 0.901 | 0.145 | 0.308 | 0.197 | 3.426 | 0.164 | 0.925 | 0.075 | 5.505 |
| CADD | 0.706 | 0.087 | 0.417 | 0.054 | 0.835 | 0.102 | 20.153 | 0.093 | 0.399 | 0.601 | 3.354 |
| Polyphen2_HDIV | 0.684 | 0.073 | 0.545 | 0.061 | 0.737 | 0.114 | 15.711 | 0.107 | 0.537 | 0.463 | 3.250 |

*Note:* <sup>a</sup>AUROC: area under the receiver operating characteristic curve; <sup>b</sup>AUPRC: area under the precision-recall curve; <sup>c</sup>MCC: Matthews correlation coefficient; <sup>d</sup>TNR: true negative rate or specificity; <sup>e</sup>FPR: false positive rate; <sup>f</sup>DOR: diagnostic odd ratio.

To further evaluate the robustness of mvPPT, we proceeded to random sampling. We repeated the random sampling for 20 rounds. In each round, 20% of the variants in the independent test set were sampled. The results showed that mvPPT displayed the highest efficiency and robustness (**Figure S5**). Since mvPPT included AFs as features, we then tested the performance of our model when lacking AF information. We compared the predictive power of mvPPT and existing methods on variants with different AF levels (based on gnomAD exomes). As shown in **Figure S6**, mvPPT performed the best on variants with different AF levels.

For further assessment, we assembled a test set with pathogenic variants from DoCM, a highly curated database of known, disease-causing mutations in cancer derived from literature, and benign variants randomly selected from VariSNP and VKGL. MvPPT again achieved the best performance in this test set (**Figure S7**).

### Performance of mvPPT on pathogenic variants within novel disease genes

To further evaluate the performance of our predictor on variants in novel disease genes, we collected novel pathogenic genes from recent publications and 62 missense variants were retained with complete scores on all comparators (see Methods and **Table S5**) (50–54). We simulated 62 exomes of Mendelian diseases, by selecting one disease-causing variant and randomly selecting 1,000 neutral variants from 1KGP. For each simulated exome, we calculated the percentage of predicted deleterious variants obtained by different predictors, according to the authors’ recommended threshold (**Figure 6A**, **Table S6**). The ranking of disease-causing variants among all variants in each simulated exome is presented in **Figure 6**. Among all predictors, PrimateAI generated the shortest the list of pathogenic variants, followed by MISTIC and mvPPT. But only 41/62 and 44/62 disease-causing variants were predicted as pathogenic by PrimateAI and MISTIC, respectively (**Table S6**). CADD identified 100% of these 62 variants as pathogenic, but may cause a plenty of false positives (PPV = 0.615 ± 0.002) (**Figure 6A**, **Table S6**). Instead, mvPPT performed relatively well in both sensitivity (60/62) and PPV (0.137 ± 0.001) (**Figure 6A**, **Table S6**). To further evaluate the ability of each predictor in prioritizing the pathogenic variants, we computed the ranking of pathogenic variants among all variants in simulated exomes. Pathogenic variants show the best ranking in mvPPT (29 ± 9), significantly better than the rest of the tools (second best, ClinPred 75 ± 11, *P*-value = 2.12E-08, Wilcoxon matched-pairs signed rank test), further demonstrating the advantages of mvPPT on detecting variants within novel disease genes.

**Figure 6.**
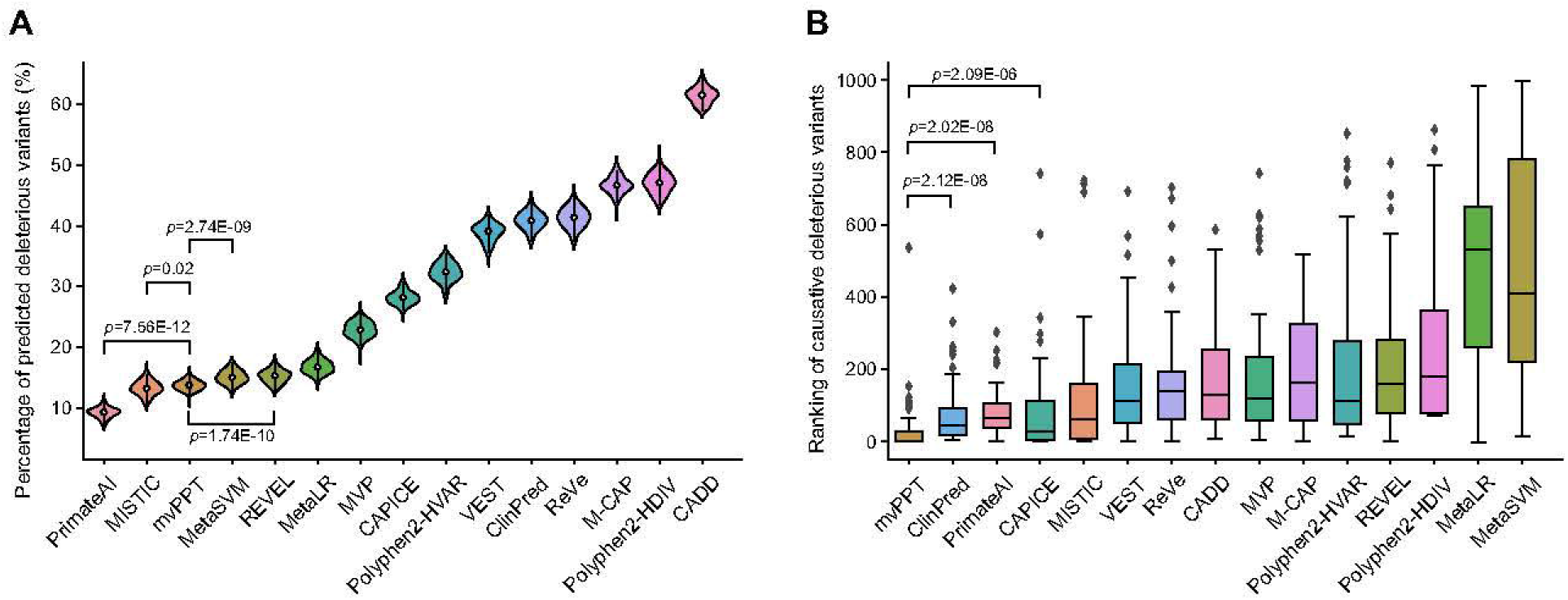
Evaluation of the different prediction tools using simulated disease exomes. (**A**) Distribution of the percentage of predicted deleterious variants in the simulated disease exomes. (**B**) Ranking of the disease-causing variants in the simulated disease exomes.

## Discussion

Missense variants as the most common category of single nucleotide variants have important implications for human genetic diseases. Although a variety of variant pathogenicity assessment tools have been established and have made important contributions to genetic variant evaluation, there is still room for improvement in prediction accuracy and precision, which is of great importance for the explanation of the tremendous number of genetic variants. In this study, we present a comprehensive prediction tool, mvPPT, and demonstrate that the performance of mvPPT is superior to all comparators in AUROC, AUPRC, F1 score and many other metrics. We found that the improvement in prediction probably resulted from careful selection of the algorithm, the features, and the training sets.

Boosting algorithms are widely adopted by many ensemble models of variant classification to improve accuracy of prediction. MvPPT adopted a recently developed LightGBM algorithm proven to outperform the existing boosting frameworks on both efficiency and accuracy. Other than the boosting algorithms, we also tried a deep neural network framework and found that our model gave higher accuracy and precision than 10-layer, 15-layer, and 20-layer fully connected neural networks (data not shown). Compared with traditional machine learning algorithms, deep learning performs better as the scale of data increases (55). We expect more accurate prediction models based on deep neural networks in the near future, with the accumulation of larger amounts of high-confidence training data. Recently developed deep learning models such as primateAI, spliceAI (56), and EVE (57) provided new perspectives of variant pathogenicity prediction, which use surrounding DNA sequences as input, without requiring explicit features. These approaches could be possibly further improved with larger training data, as well as incorporating sequence conservation, constraints, and protein structure information.

In this study, mvPPT adopted 62 features belonging to three categories. Other than commonly used features extracted from previous predictors (category A), mvPPT included two categories of features associated with allele/genotype/amino acid frequencies (category B) and genetic constraint of adjunction regions (category C). Our benchmarking studies revealed that category B and category C contributed significantly to the prediction. Among them, features in category C contributed the most as a whole, and adding category B further promotes the performance (**Figure S4**). The high predictive power of features in category B and C could be explained by the fact that natural selection constantly eliminates deleterious variants during evolution, and thus deleterious variants tend to locate in more conserved and intraspecies constrained genomic regions with lower AF/GF/AAF in human populations compared with neutral mutations. Considering that the pathogenic variants in disease databases are likely to have small AF and GF due to existing pathogenic variant selection criteria, we further tested our model on rare variants with AF = 0, and confirmed that the model works well even AF is not functioning as a feature in this case. Other than the features we used, protein structural changes corresponding to changes in amino acid sequence may also be critical predictors of variant pathogenicity. Newly developed protein structure prediction tools, such as AlphaFold2 (58,59) have made it possible to include protein structure information in future tools.

In addition to the algorithm and features, the improvement in performance of mvPPT also came from a cautious selection of the training set. While most of the prevailing variant classifiers trained their models on single databases, previous studies have uncovered considerable disagreement among databases (7,9). Some tools also enrolled variants in general population databases (e.g., Exome Aggregation Consortium (60)) as benign variants, to increase the size of the training set. But this setting may add noise to the training data, as not all variants in population databases are truly benign. In this study, we found unlike adding full sets of variants in gnomAD, which attenuates the performance of the prediction model, adding variants from with AF above a certain threshold in general enhances the model. The possible explanation of this observation is that the singletons are more likely to be contaminated with false benign variants or less confident variants compared with non-singletons. Therefore, we examined the labels of singletons from gnomAD that we added to the training set in ClinVar. Among gnomAD with labels in ClinVar, we found that 8.1% of them were benign (labeled as ‘Benign’, ‘Likely benign’, and ‘Benign/Likely benign’), 78.5% were labeled as ‘uncertain’, 7.4% were labeled as ‘Conflicting interpretations of pathogenicity’, and 6.0% were pathogenic (labeled as ‘Pathogenic’, ‘Likely pathogenic’, and ‘Pathogenic/Likely pathogenic’). In contrast, among the overlapped non-singleton missense variants, 47.8% were benign, 34.8% were labeled as ‘uncertain’, 15.3% were labeled as ‘Conflicting interpretations of pathogenicity’, and 2.1% were pathogenic. Overall, our observation highlights the importance of maintaining a balance between size and purity of the training set, as well as, provides a practical guidance of training set selection.

As reported by previous studies, we observed that the pathogenic variants in disease databases are enriched in certain genes. However, our down-sampling experiments suggest that the aggregation of variants has few effects on pathogenicity prediction. This can be possibly explained by the fact that most features of variants are independent of genes. Furthermore, we found that adding region/gene-based information (category C) slowed down the effects of down-sampling, suggesting that incorporating genomic context further lessens the impacts the variants aggregation (**Figures S2B**, **C**).

In conclusion, we developed an ensemble classifier, mvPPT, for predicting the pathogenicity of missense variants, and demonstrated that mvPPT achieved superior performance compared with other established prediction tools. Particularly, in clinical data, mvPPT showed the highest accuracy and robustness in classifying variants associated with both Mendelian diseases and cancer. Therefore, mvPPT promises to facilitate a better clinical interpretation of missense variants with uncertain significance. For convenient use, we built a searchable website and all pre-computed mvPPT scores are available at http://www.mvppt.club. All codes are available at https://github.com/tongshiyuan/mvPPT.

## Materials and methods

Web resources of all databases and software used in the manuscript are listed in **Table S7**.

### Missense variant annotation

Variants were annotated by the latest version of ANNOVAR software (version 2019Oct24) (61), with gene-based annotation set to ensSeq (assembly version hg19). Variants whose functional consequences were marked as nonsynonymous SNV were selected. Further removing loss-of-function (stop gain or stop loss) variants ensured that our model was trained and evaluated nearly exclusively on missense variants.

### Training set

Training set variants were collected from disease databases: ClinVar (2020.07), HGMD (Pro version 2020.3), and UniProt (2020.06), as well as population database gnomAD genomes (version 2.1.1, see Web Resources). Each variant in ClinVar has a review status tag reporting the level of review supporting the assertion of clinical significance, and a clinical significance tag labeling variants as pathogenic, likely pathogenic, uncertain significance, likely benign, and benign. To select variants with reliable tags, we kept variants with review status of “criteria provided” from submitters and “reviewed by expert panel”. The variants were further filtered according to their significance tag: variants that were categorized as (1) benign or likely benign and (2) pathogenic or likely pathogenic were selected as negative (benign) and positive (pathogenic) labels, respectively. The variants in HGMD were labeled by 7 different tags, including DM (disease-causing mutation), DM? (disease-causing mutation?), DP (disease-associated polymorphism), DFP (disease-associated polymorphism with supporting functional evidence), FP (in vitro/laboratory or in vivo functional polymorphism), FTV (polymorphic or rare variants reported in the literature), and R (Retired entry). DM and DM? label variants reported to be disease causing in the original literature report. The question mark denotes that a degree of doubt has been found with regard to pathogenicity. We only kept the variants with “DM” label in this study. For UniProt, there are three variant labels: Disease, Polymorphism, and Unclassified. These labels were curated from literature reports. We kept the variants labeled with “Disease” and “Polymorphism” in this study. All variants with conflict label in different databases were excluded. Population variants were obtained from gnomAD genomes (version 2.1.1), which combines variation data from 15,708 individuals. To avoid any bias, the population variants were further filtered to remove any variants in disease databases. Ten-fold cross-validation was implemented through the Python package scikit-learn version 0.23.2.

### Cross-validation in algorithm selection and feature selection

To avoid overfitting, we designed the ten-fold cross-validation procedure as follows: 1) we divided the variants from disease and 1KGP databases into ten subsets; 2) in each round, we selected nine subsets of variants from disease databases to generate the training set, and combined the rest one subset of variants from disease databases with one subset of 1KGP variants to form the test set.

### Cross-validation in training set analysis

The same cross-validation procedure was conducted as above, except that in each round, the training set was generated using variants from disease databases only, or adding variants from gnomAD as benign variants. More specifically, we first divided the variants from disease databases into 10 subsets. In each round, we selected 90% of variants from the disease databases and combined them with variants in gnomAD genomes passing different AF thresholds (all, removing singleton, AF > 0.001, AF > 0.001, and AF > 0.01) to generate the training sets for comparison; the rest 10% of variants from disease databases were then combined with 10% variants from 1000 Genomes to form the test set.

### Test set

Test set combined variants from VKGL (September 2020), VariSNP (2017-02-16), DoCM (version 3.2), DPV (2020.12.29), CAPICE_Test (v4), and MetaLR/SVM_Test. To guarantee a second independent test set with variants that has never been used in any tools’ training set, we used PubMed to search for papers reporting new genetic disease-causing genes. In total, we found 5 papers, covering 7 genes, and none of these genes include pathogenic variants in the training or the test sets in our study. All missense variants on these genes reported in the literatures were collected. Population variants were obtained from 1KGP. To validate the robustness of our model and avoid overfitting, variants that were used in any training set or in our features’ training data were discarded from all test sets, and only variants with comprehensive scores required by all comparator models were included.

### Features

MvPPT adopted 62 features from three categories: A) pathogenicity likelihood scores assessed by different component tools, including MutationAssessor, SIFT, PROVEAN, GERP++RS, phyloP, phastCons, and SiPhy. B) AFs, GFs, and AAFs of variants estimated from 125,748 exomes in gnomAD (version 2.1.1); C) genomic context of the variant, i.e., region/gene-based information from GeVIR, VIRLoF, oe_mis_upper (from gnomAD), HIP, CCRs, Interpro domain, and amino acid sequence before and after mutation. To avoid overfitting, the 7 tools we used in category A did not generate scores based on machine learning algorithms. We annotated datasets with ANNOVAR using dbNSFP database (v.4.1a) (62,63) to generate some of the required prediction scores from different component tools, including Interpro domain, MutationAssessor, phyloP, GERP, phastCons, PROVEAN, and SiPhy. Mutations located in the interpro domains were recorded as 1 and the rest were recorded as 0. AFs, GFs, and AAFs of each variant in different populations were obtained from the gnomAD exomes. AFs, AAFs, HomFs, and HetFs were assigned 0 and WtFs were assigned 1 if the variant was not present in the database. The GeVIR, VIRLoF, oe_mis_upper, HIP, and CCR scores were downloaded from their respective websites. One-hot encoding has been applied to amino acid sequence, representing each amino acid with a binary vector of length 20 with a single non-zero value. All the features were selected to provide complementary information, and they either did not require training or their training data are publicly available to allow exclusion from our data.

### Outlier detection and GO enrichment analysis

The interquartile range (IQR) was used to identify outliers. The IQR criterion is summarized as follows:

1. Compute the first and third quartiles, Q_1*j*_ and Q_3*j*_, for each peptide *j*, and then its IQR: IQR_*j*_ = Q_3*j*_ – Q_1*j*_.
2. For each peptide *j*, observation *y_ij_* is flagged as an outlier if *y_ij_* < Q_1*j*_ – k IQR_*j*_ or *y_ij_* > Q_3*j*_ + k IQR_*j*_, where k = 1.5.

GO enrichment analysis was performed with the R package clusterProfiler (64).

### Metrics for performance evaluation

We used 11 different metrics to evaluate the performance of the prediction tools. Detailed description of the metrics was provided in **Table S8**.

### MvPPT training

MvPPT was trained using the python package LightGBM (version 2.3.1) (31), and parameters were tuned by Bayesian optimization (version 1.2.0). The random status was set as 1 throughout the model training process. For Bayesian optimization process, the number of iterations was set as 100 (n_iter = 100) and the steps of random exploration was set as 15 (init_points = 15). The ranges of the hyperparameters in the LightGBM for Bayesian optimization were set as follows: num_leaves (24, 45), feature_fraction (0.1, 0.9), bagging_fraction (0.8, 1), max_depth (5, 8.99), lambda_l1 (0, 5), lambda_l2 (0, 3), min_split_gain (0.001, 0.1), min_child_weight (5, 50). After the parameter optimization process, the final used value of the parameters was as follows: num_leaves = 45, min_child_weight = 6.163, learning_rate = 0.01, bagging_fraction = 0.870, feature_fraction = 0.632, lambda_l1 = 0. 921, lambda_l2 = 0.193, min_gain_to_split = 0.039, and max_depth = 9.

### Scores from existing tools

The MVP (12), REVEL (6), PrimateAI (8), FATHMM-XF (20), ClinPred (7), MetaSVM/MetaLR (9), PolyPhen2 (16), and VEST4 (11) scores were obtained from dbNSFP v4.1a. The MISTIC (13), CAPICE (19), M-CAP (5), ReVe (22), and CADD (21) scores were downloaded from their respective websites.

### Statistical analysis

Wilcoxon matched-pairs signed rank test was conducted using the stats module in SciPy Python package version 1.5.4. *P*.adjust in GO enrichment analysis was calculated by the R package clusterProfiler. All the metrics in this paper were calculated based on the scikit-learn Python package.

### Data and code availability

The mvPPT scores for potential missense variants in the human genome are available at mvPPT website (**Table S7**). The mvPPT code is available at GitHub (**Table S7**) for noncommercial purposes.

## Authors’ contributions

Y.-C.Y. and Y.Z. conceived the analysis. S.-Y.T., K.F., and Z.-W.Z. collected the databases. S.-Y.T. and Z.-W.Z. performed the analysis. S.-Y.T., Y.Z., and Y.-C.Y. drafted the manuscript. K.F., L.-Y.L., S.-Q.Z., and Y.F. rechecked the results. S.-Y.T. and G.-Z.W. built the website. All authors read and approved the final manuscript.

## Acknowledgments

This work is supported by the Shanghai Natural Science Foundation [grant numbers 20ZR1403800]; the National Natural Science Foundation of China [grant numbers 31900476, 82071259, 31930044, 31725012]; the Shanghai Municipal Science and Technology Major Project [grant numbers 2018SHZDZX01] and ZJLab; the Foundation of Shanghai Municipal Education Commission [grant numbers 2019-01-07-00-07-E00062]; and the Collaborative Innovation Program of Shanghai Municipal Health Commission [grant numbers 2020CXJQ01].

## Supplementary materials

Figures

**Figure S1 The prediction power of features by category**

(**A**) The AUROC and (**B**) the AUPRC obtained from models trained on each combination of categories were plotted. We excluded 1KGP variants from the test set in **Figure 2** and re-conducted the comparisons.

**Figure S2 Down-sampling**

(**A**) Thresholds for down-sampling. (**B**) AUROC and (**C**) AUPRC obtained from models trained on variants selected according to different down-sampling thresholds were plotted.

**Figure S3 Allele frequency distribution of pathogenic/begin variants in different data sets**

The x axis shows the allele frequency based on gnomAD exomes (For convenience, only the variants with AF < 0.01 are shown). The y axis denotes the relative frequency. (**A**) One randomly selected control training set of the 10-fold training data sets in the study. (**B**) The gnomAD genomes variants with AF > 0.01 were used as benign variants and added to **A**. (**C**) The gnomAD genomes variants with AF > 0.001 were used as benign variants and added to **A**. (**D**) the gnomAD genomes variants with AF > 0.0001 were used as benign variants and added to **A**. (**E**) The gnomAD genomes variants that removed singleton variants were used as benign variants and added to **A**. (**F**) All of the gnomAD genomes variants were used as benign variants and added to **A**. (**G**) The paired test set data corresponding to the training set data. (**H**) The collection of all the variants in the HGMD, ClinVar, and UniProt.

**Figure S4 The correlation among the individual and feature importance in mvPPT**

(**A**) Correlation among the individual features, ordered by hierarchical clustering. The heatmap illustrates the Spearman rank correlation coefficients between features computed for the mvPPT training variants. (**B**) An overview of the relative importance of the structured features integrated. Relative importance was calculated by normalizing Gini importance to sum to one.

**Figure S5 Comparing the performance of different prediction tools by random sampling**

(**A**) The AUROC, (**B**) the AUPRC and (**C**) the efficiency of mvPPT and 15 established prediction methods when tested on an independent test set were plotted. This random sampling was repeated 20 iterations. In each iteration, 20% of variants were sampled. Efficiency was obtained by calculating the mean of sensitivity, specificity, and accuracy.

**Figure S6 The performance of predictors stratified by allele frequency (AF)**

(**A**) The AUROC and (**B**) the AUPRC of mvPPT and 15 established prediction methods for independent test set stratified by variant AF.

**Figure S7 Comparing the performance of mvPPT on cancer test set**

(**A**) AUROC and (**B**) AUPRC of mvPPT and 15 established prediction methods when tested on cancer test set (DoCM, VariSNP, and VKGL) were plotted.

Tables

**Table S1 List of outlier genes**

**Table S2 Overview of performance of models with different training set**

**Table S3 Description of training set**

**Table S4 List of test set**

**Table S5 List of missense variants in novel genes**

**Table S6 List of thresholds used for the predictors and variants predicted to be pathogenic**

**Table S7 Description of databases and softwares**

**Table S8 Description about the metrics used to evaluate the performance of the prediction tools**

## References

1. Lee H, Deignan JL, Dorrani N, Strom SP, Kantarci S, Quintero-Rivera F, et al. Clinical Exome Sequencing for Genetic Identification of Rare Mendelian Disorders. JAMA. 2014 Nov 12;312(18):1880.

2. Yang Y, Muzny DM, Reid JG, Bainbridge MN, Willis A, Ward PA, et al. Clinical Whole-Exome Sequencing for the Diagnosis of Mendelian Disorders. N Engl J Med. 2013 Oct 17;369(16):1502–11.

3. Shihab HA, Gough J, Mort M, Cooper DN, Day IN, Gaunt TR. Ranking non-synonymous single nucleotide polymorphisms based on disease concepts. Hum Genomics. 2014;8(1):11.

4. Ng PC, Levy S, Huang J, Stockwell TB, Walenz BP, Li K, et al. Genetic Variation in an Individual Human Exome. Schork NJ, editor. PLoS Genet. 2008 Aug 15;4(8):e1000160.

5. Jagadeesh KA, Wenger AM, Berger MJ, Guturu H, Stenson PD, Cooper DN, et al. M-CAP eliminates a majority of variants of uncertain significance in clinical exomes at high sensitivity. Nature Genetics. 2016 Dec;48(12):1581–6.

6. Ioannidis NM, Rothstein JH, Pejaver V, Middha S, McDonnell SK, Baheti S, et al. REVEL: An Ensemble Method for Predicting the Pathogenicity of Rare Missense Variants. The American Journal of Human Genetics. 2016 Oct;99(4):877–85.

7. Alirezaie N, Kernohan KD, Hartley T, Majewski J, Hocking TD. ClinPred: Prediction Tool to Identify Disease-Relevant Nonsynonymous Single-Nucleotide Variants. The American Journal of Human Genetics. 2018 Oct 4;103(4):474–83.

8. Sundaram L, Gao H, Padigepati SR, McRae JF, Li Y, Kosmicki JA, et al. Predicting the clinical impact of human mutation with deep neural networks. Nature Genetics. 2018 Aug;50(8):1161–70.

9. Dong C, Wei P, Jian X, Gibbs R, Boerwinkle E, Wang K, et al. Comparison and integration of deleteriousness prediction methods for nonsynonymous SNVs in whole exome sequencing studies. Human Molecular Genetics. 2015 Apr 15;24(8):2125–37.

10. Kumar P, Henikoff S, Ng PC. Predicting the effects of coding non-synonymous variants on protein function using the SIFT algorithm. Nature Protocols. 2009 Jul;4(7):1073–81.

11. Carter H, Douville C, Stenson PD, Cooper DN, Karchin R. Identifying Mendelian disease genes with the Variant Effect Scoring Tool. BMC Genomics. 2013;14(Suppl 3):S3.

12. Qi H, Zhang H, Zhao Y, Chen C, Long JJ, Chung WK, et al. MVP predicts the pathogenicity of missense variants by deep learning. Nat Commun. 2021 Dec;12(1):510.

13. Chennen K, Weber T, Lornage X, Kress A, Böhm J, Thompson J, et al. MISTIC: A prediction tool to reveal disease-relevant deleterious missense variants. Andrade-Navarro MA, editor. PLoS ONE. 2020 Jul 31;15(7):e0236962.

14. Ip E, Chapman G, Winlaw D, Dunwoodie SL, Giannoulatou E. VPOT: A Customizable Variant Prioritization Ordering Tool for Annotated Variants. Genomics, Proteomics & Bioinformatics. 2019 Oct;17(5):540–5.

15. Reva B, Antipin Y, Sander C. Predicting the functional impact of protein mutations: application to cancer genomics. Nucleic Acids Research. 2011 Sep;39(17):e118–e118.

16. Adzhubei IA, Schmidt S, Peshkin L, Ramensky VE, Gerasimova A, Bork P, et al. A method and server for predicting damaging missense mutations. Nature Methods. 2010 Apr;7(4):248–9.

17. Schwarz JM, Cooper DN, Schuelke M, Seelow D. MutationTaster2: mutation prediction for the deep-sequencing age. Nature Methods. 2014 Apr;11(4):361–2.

18. Li Q, Liu X, Gibbs RA, Boerwinkle E, Polychronakos C, Qu H-Q. Gene-Specific Function Prediction for Non-Synonymous Mutations in Monogenic Diabetes Genes. Brusgaard K, editor. PLoS ONE. 2014 Aug 19;9(8):e104452.

19. Li S, van der Velde KJ, de Ridder D, van Dijk ADJ, Soudis D, Zwerwer LR, et al. CAPICE: a computational method for Consequence-Agnostic Pathogenicity Interpretation of Clinical Exome variations. Genome Med. 2020 Dec;12(1):75.

20. Rogers MF, Shihab HA, Mort M, Cooper DN, Gaunt TR, Campbell C. FATHMM-XF: accurate prediction of pathogenic point mutations via extended features. Bioinformatics. 2018 Feb 1;34(3):511–3.

21. Rentzsch P, Witten D, Cooper GM, Shendure J, Kircher M. CADD: predicting the deleteriousness of variants throughout the human genome. Nucleic Acids Research. 2019 Jan 8;47(D1):D886–94.

22. Li J, Zhao T, Zhang Y, Zhang K, Shi L, Chen Y, et al. Performance evaluation of pathogenicity-computation methods for missense variants. Nucleic Acids Res. 2018 Sep 6;46(15):7793–804.

23. Landrum MJ, Lee JM, Benson M, Brown GR, Chao C, Chitipiralla S, et al. ClinVar: improving access to variant interpretations and supporting evidence. Nucleic Acids Research. 2018 Jan 4;46(D1):D1062–7.

24. Stenson PD, Mort M, Ball EV, Evans K, Hayden M, Heywood S, et al. The Human Gene Mutation Database: towards a comprehensive repository of inherited mutation data for medical research, genetic diagnosis and next-generation sequencing studies. Hum Genet. 2017 Jun;136(6):665–77.

25. Peterson TA, Doughty E, Kann MG. Towards Precision Medicine: Advances in Computational Approaches for the Analysis of Human Variants. Journal of Molecular Biology. 2013 Nov 1;425(21):4047–63.

26. Salnikova LE, Kolobkov DS, Sviridova DA, Abilev SK. An overview of germline variations in genes of primary immunodeficiences through integrative analysis of ClinVar, HGMD^®^ and dbSNP databases. Hum Genet. 2021 Sep;140(9): 1379–93.

27. Tennessen JA, Bigham AW, O’Connor TD, Fu W, Kenny EE, Gravel S, et al. Evolution and Functional Impact of Rare Coding Variation from Deep Sequencing of Human Exomes. Science. 2012 Jul 6;337(6090):64–9.

28. Genome Aggregation Database Consortium, Karczewski KJ, Francioli LC, Tiao G, Cummings BB, Alföldi J, et al. The mutational constraint spectrum quantified from variation in 141,456 humans. Nature. 2020 May;581(7809):434–43.

29. Abramovs N, Brass A, Tassabehji M. GeVIR is a continuous gene-level metric that uses variant distribution patterns to prioritize disease candidate genes. Nat Genet. 2020 Jan;52(1):35–9.

30. Vitsios D, Dhindsa RS, Middleton L, Gussow AB, Petrovski S. Prioritizing non-coding regions based on human genomic constraint and sequence context with deep learning. Nat Commun. 2021 Dec;12(1):1504.

31. Ke G, Meng Q, Finley T, Wang T, Chen W, Ma W, et al. LightGBM: A Highly Efficient Gradient Boosting Decision Tree. In: Guyon I, Luxburg UV, Bengio S, Wallach H, Fergus R, Vishwanathan S, et al., editors. Advances in Neural Information Processing Systems 30 [Internet]. Curran Associates, Inc.; 2017 [cited 2020 Jul 7]. p. 3146–54. Available from: http://papers.nips.cc/paper/6907-lightgbm-a-highly-efficient-gradient-boosting-decision-tree.pdf

32. Anghel A, Papandreou N, Parnell T, De Palma A, Pozidis H. Benchmarking and Optimization of Gradient Boosting Decision Tree Algorithms. arXiv:180904559 [cs, stat] [Internet]. 2019 Jan 17 [cited 2020 Sep 22]; Available from: http://arxiv.org/abs/1809.04559

33. The UniProt Consortium. UniProt: a worldwide hub of protein knowledge. Nucleic Acids Research. 2019 Jan 8;47(D1):D506–15.

34. The 1000 Genomes Project Consortium. A global reference for human genetic variation. Nature. 2015 Oct;526(7571):68–74.

35. Pollard KS, Hubisz MJ, Rosenbloom KR, Siepel A. Detection of nonneutral substitution rates on mammalian phylogenies. Genome Research. 2010 Jan 1;20(1):110–21.

36. Davydov EV, Goode DL, Sirota M, Cooper GM, Sidow A, Batzoglou S. Identifying a High Fraction of the Human Genome to be under Selective Constraint Using GERP++. Wasserman WW, editor. PLoS Computational Biology. 2010 Dec 2;6(12):e1001025.

37. Siepel A. Evolutionarily conserved elements in vertebrate, insect, worm, and yeast genomes. Genome Research. 2005 Aug 1;15(8):1034–50.

38. Choi Y, Sims GE, Murphy S, Miller JR, Chan AP. Predicting the Functional Effect of Amino Acid Substitutions and Indels. de Brevern AG, editor. PLoS ONE. 2012 Oct 8;7(10):e46688.

39. Garber M, Guttman M, Clamp M, Zody MC, Friedman N, Xie X. Identifying novel constrained elements by exploiting biased substitution patterns. Bioinformatics. 2009 Jun 15;25(12):i54–62.

40. Huang N, Lee I, Marcotte EM, Hurles ME. Characterising and Predicting Haploinsufficiency in the Human Genome. Schierup MH, editor. PLoS Genet. 2010 Oct 14;6(10):e1001154.

41. Havrilla JM, Pedersen BS, Layer RM, Quinlan AR. A map of constrained coding regions in the human genome. Nat Genet. 2019 Jan;51(1):88–95.

42. Jones P, Binns D, Chang H-Y, Fraser M, Li W, McAnulla C, et al. InterProScan 5: genome-scale protein function classification. Bioinformatics. 2014 May 1;30(9):1236–40.

43. Ashburner M, Ball CA, Blake JA, Botstein D, Butler H, Cherry JM, et al. Gene Ontology: tool for the unification of biology. Nat Genet. 2000 May;25(1):25–9.

44. Mi H, Muruganujan A, Ebert D, Huang X, Thomas PD. PANTHER version 14: more genomes, a new PANTHER GO-slim and improvements in enrichment analysis tools. Nucleic Acids Research. 2019 Jan 8;47(D1):D419–26.

45. The Gene Ontology Consortium. The Gene Ontology resource: enriching a GOld mine. Nucleic Acids Research. 2021 Jan 8;49(D1):D325–34.

46. Schaafsma GCP, Vihinen M. VariSNP, A Benchmark Database for Variations From dbSNP. Human Mutation. 2015 Feb;36(2):161–6.

47. Fokkema IFAC, Velde KJ, Slofstra MK, Ruivenkamp CAL, Vogel MJ, Pfundt R, et al. Dutch genome diagnostic laboratories accelerated and improved variant interpretation and increased accuracy by sharing data. Human Mutation. 2019 Dec;40(12):2230–8.

48. Suzuki H, Kurosawa K, Fukuda K, Ijima K, Sumazaki R, Saito S, et al. Japanese pathogenic variant database: DPV. Translational Science of Rare Diseases. 2018 Jul 18;1–5.

49. Ainscough BJ, Griffith M, Coffman AC, Wagner AH, Kunisaki J, Choudhary MN, et al. DoCM: a database of curated mutations in cancer. Nat Methods. 2016 Oct;13(10):806–7.

50. Fliedner A, Kirchner P, Wiesener A, van de Beek I, Waisfisz Q, van Haelst M, et al. Variants in SCAF4 Cause a Neurodevelopmental Disorder and Are Associated with Impaired mRNA Processing. The American Journal of Human Genetics. 2020 Sep;107(3):544–54.

51. Palencia-Campos A, Aoto PC, Machal EMF, Rivera-Barahona A, Soto-Bielicka P, Bertinetti D, et al. Germline and Mosaic Variants in PRKACA and PRKACB Cause a Multiple Congenital Malformation Syndrome. The American Journal of Human Genetics. 2020 Nov;107(5):977–88.

52. Tsai M-H, Muir AM, Wang W-J, Kang Y-N, Yang K-C, Chao N-H, et al. Pathogenic Variants in CEP85L Cause Sporadic and Familial Posterior Predominant Lissencephaly. Neuron. 2020 Apr;106(2):237–245.e8.

53. Hadjadj J, Castro CN, Tusseau M, Stolzenberg M-C, Mazerolles F, Aladjidi N, et al. Early-onset autoimmunity associated with SOCS1 haploinsufficiency. Nat Commun. 2020 Dec;11(1):5341.

54. Lessel D. Germline AGO2 mutations impair RNA interference and human neurological development.:14.

55. Cao C, Liu F, Tan H, Song D, Shu W, Li W, et al. Deep Learning and Its Applications in Biomedicine. Genomics, Proteomics & Bioinformatics. 2018 Feb;16(1):17–32.

56. Jaganathan K, Kyriazopoulou Panagiotopoulou S, McRae JF, Darbandi SF, Knowles D, Li YI, et al. Predicting Splicing from Primary Sequence with Deep Learning. Cell. 2019 Jan;176(3):535–548.e24.

57. Frazer J, Notin P, Dias M, Gomez A, Min JK, Brock K, et al. Disease variant prediction with deep generative models of evolutionary data. Nature. 2021 Nov 4;599(7883):91–5.

58. Jumper J, Evans R, Pritzel A, Green T, Figurnov M, Ronneberger O, et al. Highly accurate protein structure prediction with AlphaFold. Nature. 2021 Aug 26;596(7873):583–9.

59. Varadi M, Anyango S, Deshpande M, Nair S, Natassia C, Yordanova G, et al. AlphaFold Protein Structure Database: massively expanding the structural coverage of protein-sequence space with high-accuracy models. Nucleic Acids Research. 2021 Nov 17;gkab1061.

60. Exome Aggregation Consortium, Lek M, Karczewski KJ, Minikel EV, Samocha KE, Banks E, et al. Analysis of protein-coding genetic variation in 60,706 humans. Nature. 2016 Aug;536(7616):285–91.

61. Wang K, Li M, Hakonarson H. ANNOVAR: functional annotation of genetic variants from high-throughput sequencing data. Nucleic Acids Research. 2010 Sep 1;38(16):e164–e164.

62. Liu X, Jian X, Boerwinkle E. dbNSFP: A lightweight database of human nonsynonymous SNPs and their functional predictions. Hum Mutat. 2011 Aug;32(8): 894–9.

63. Liu X. dbNSFP v4: a comprehensive database of transcript-specific functional predictions and annotations for human nonsynonymous and splice-site SNVs. 2020;8.

64. Yu G, Wang L-G, Han Y, He Q-Y. clusterProfiler: an R Package for Comparing Biological Themes Among Gene Clusters. OMICS: A Journal of Integrative Biology. 2012 May;16(5):284–7.

