## Supplementary figures and images for "MvPPT: a highly efficient and sensitive pathogenicity prediction tool for missense variants"

### Supplemental Figure 1

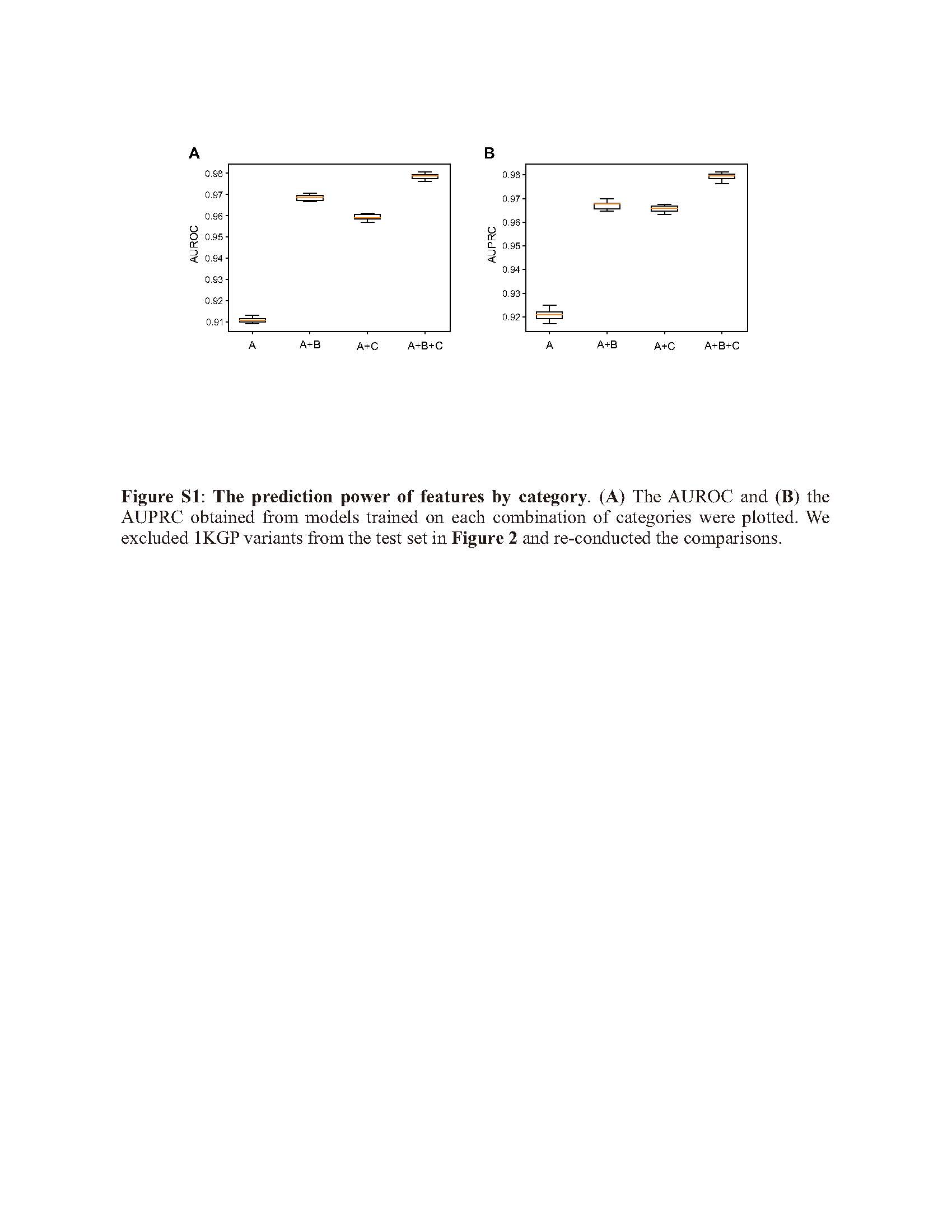

### Supplemental Figure 2

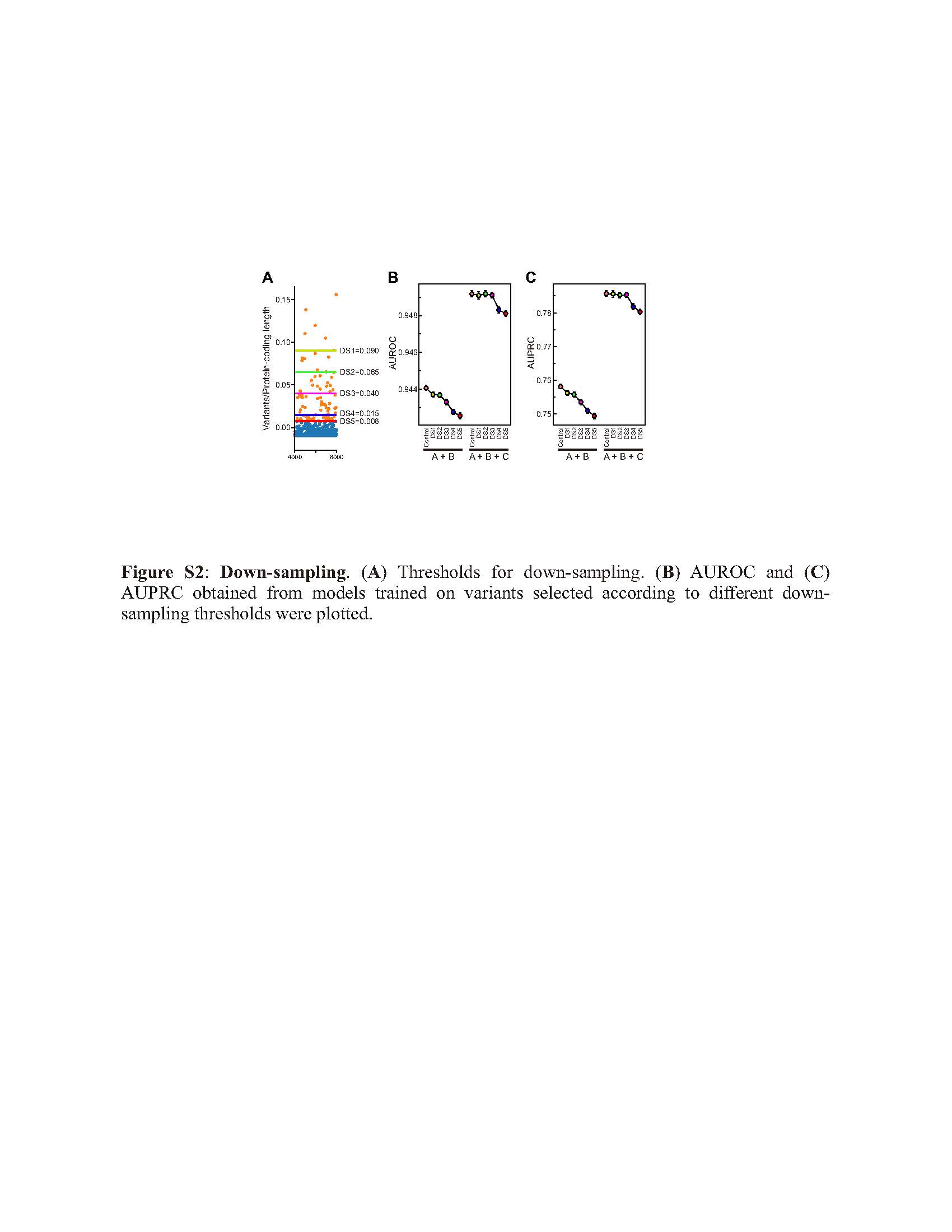

### Supplemental Figure 3

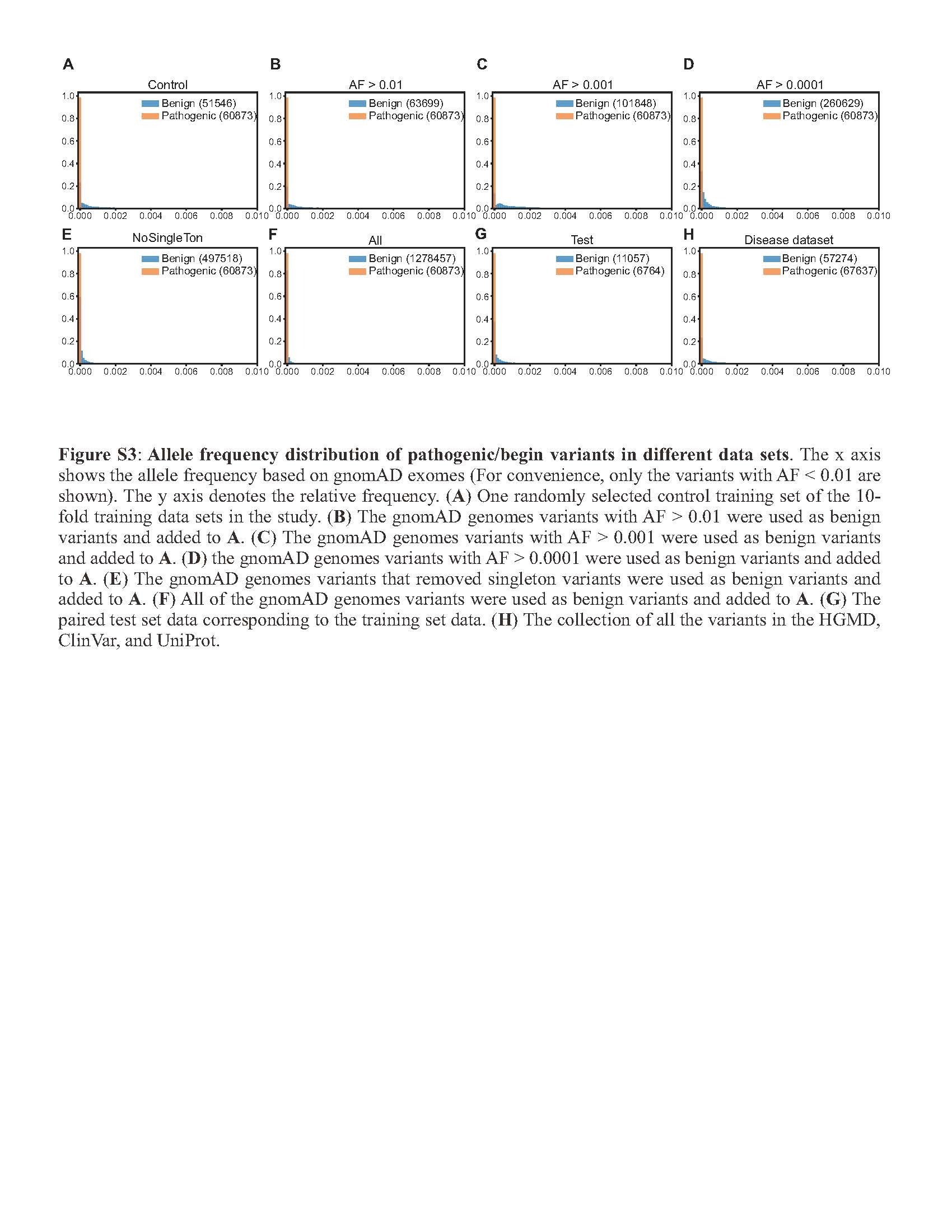

### Supplemental Figure 4

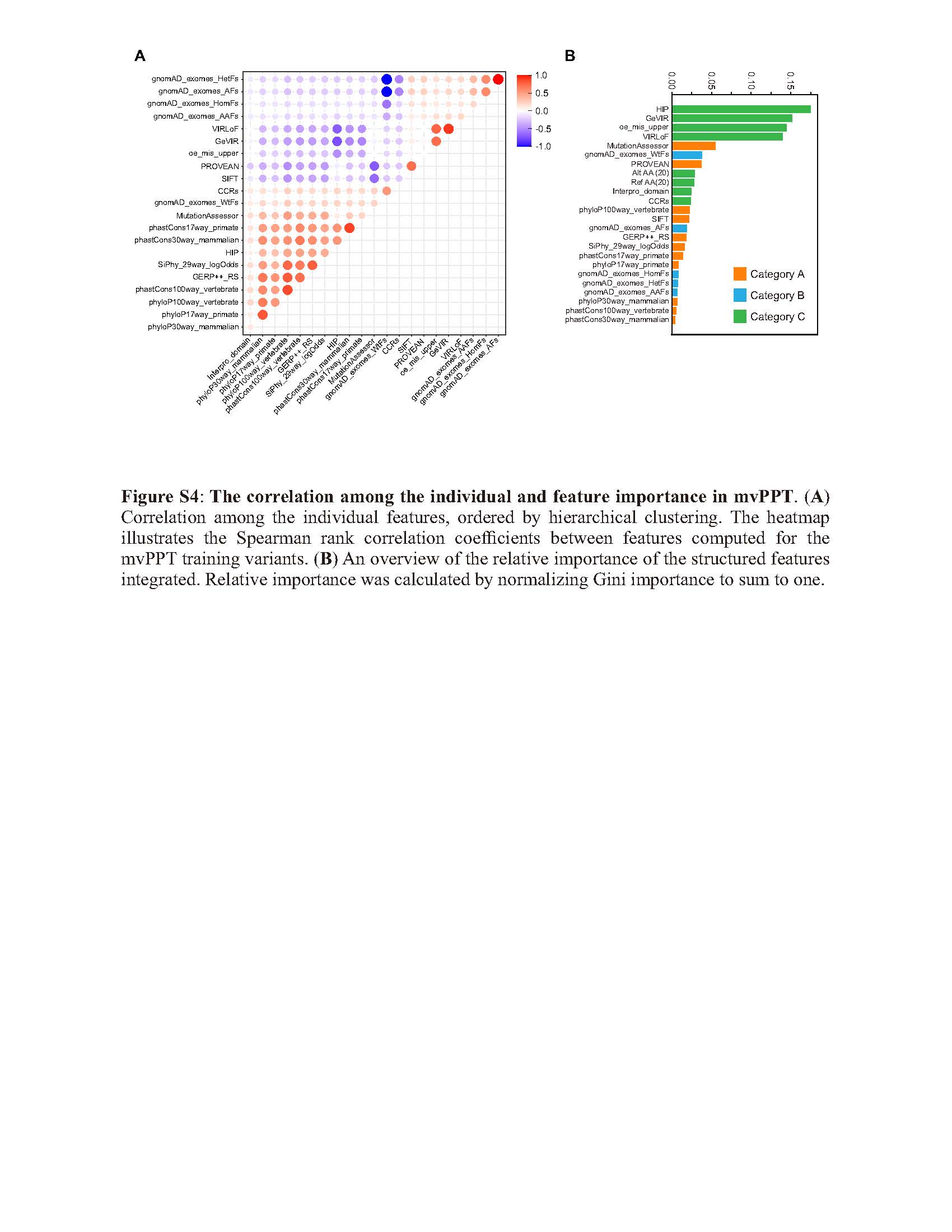

### Supplemental Figure 5

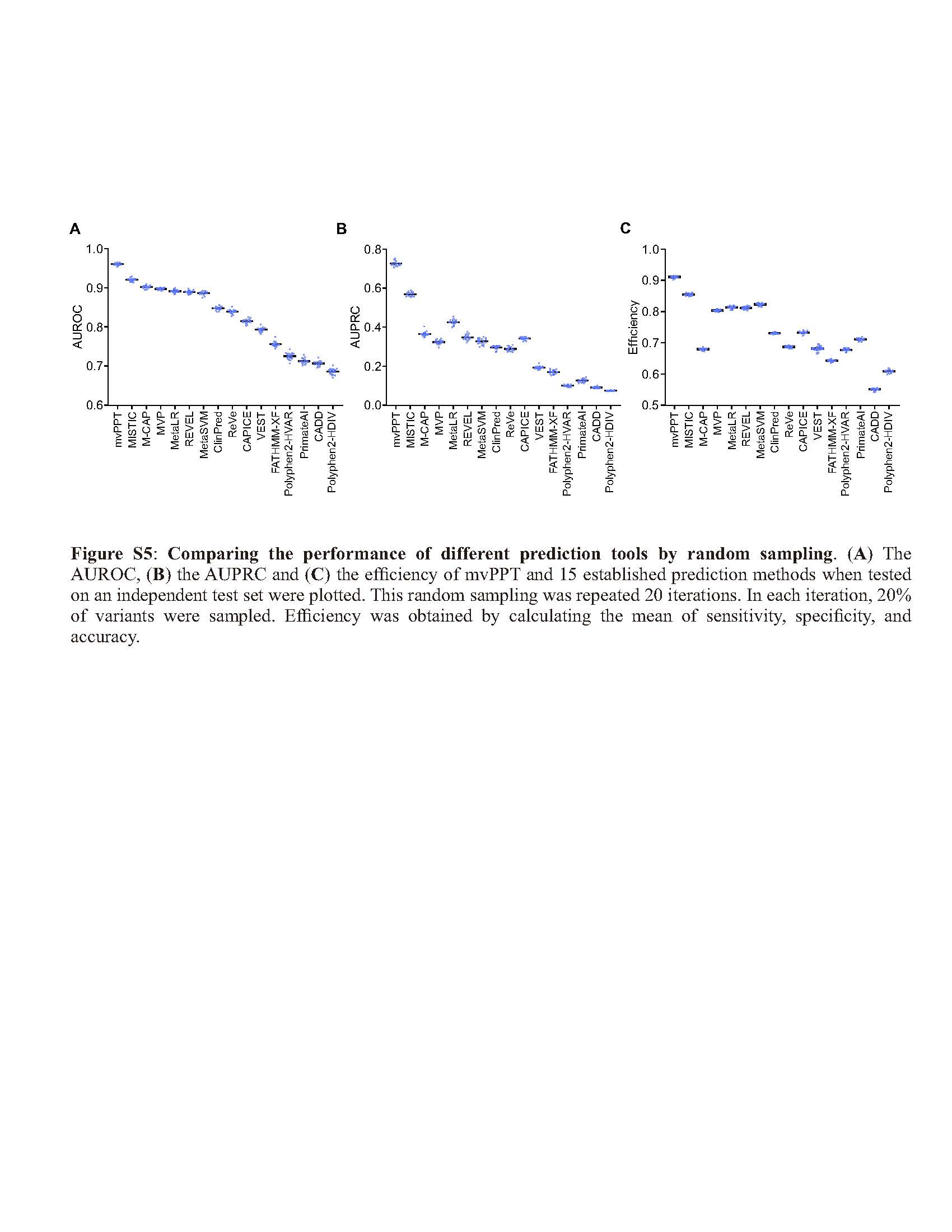

### Supplemental Figure 6

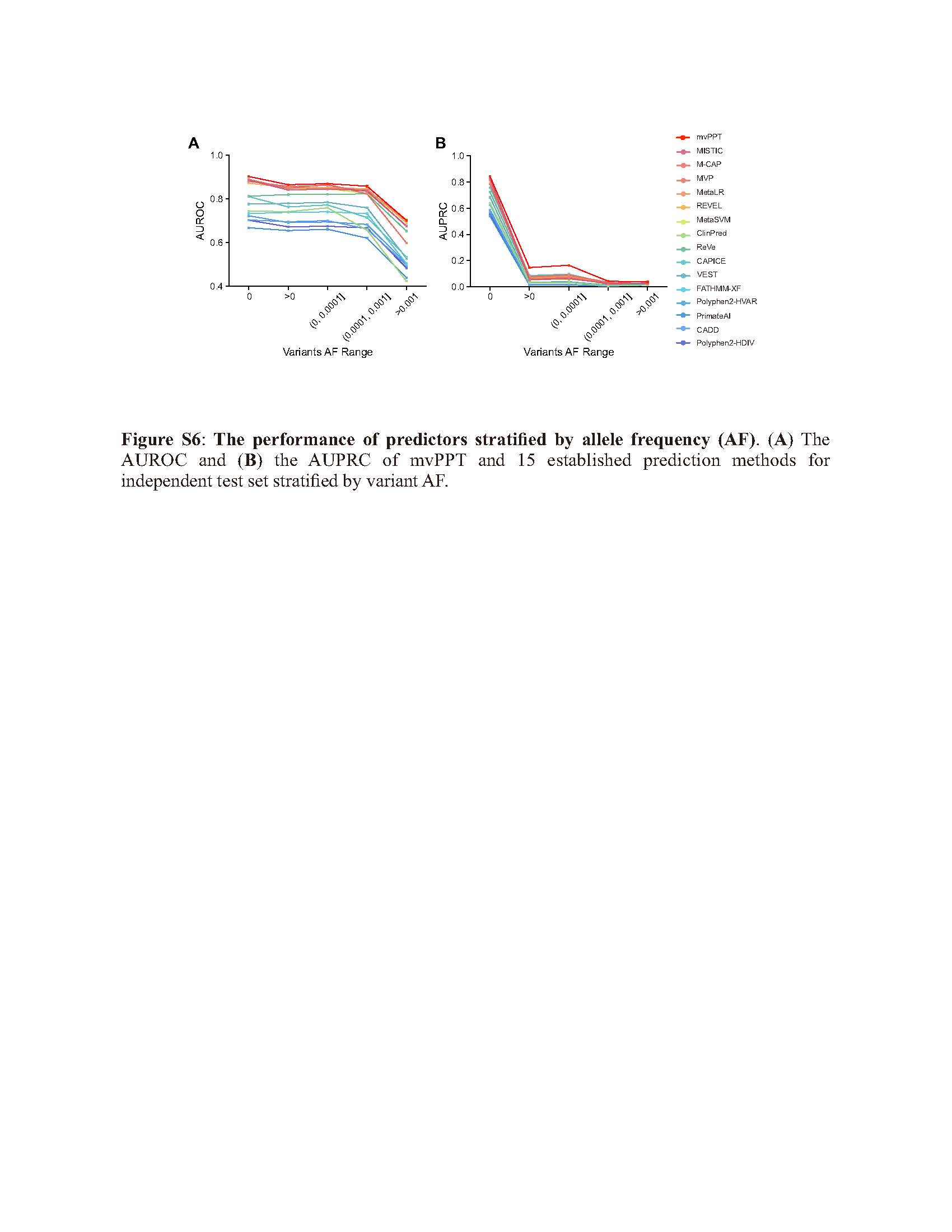

### Supplemental Figure 7

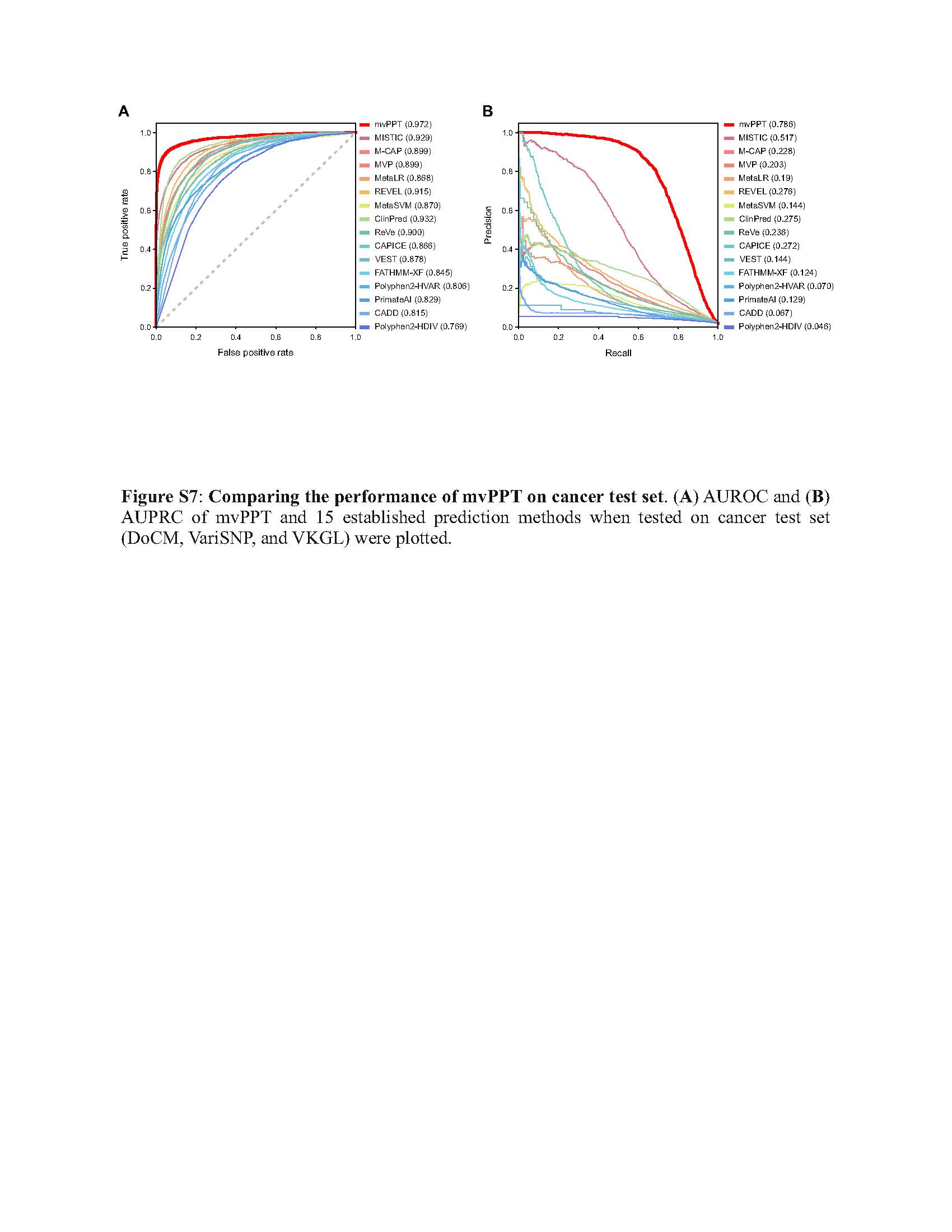
